# Monarch butterfly trends reported in Boyle et al. (2019) are sensitive to unexamined changes in museum collections over time

**DOI:** 10.1101/562314

**Authors:** Tyson Wepprich

## Abstract

Museum records can document long-term changes in phenology, species interactions, and trait evolution. However, these data have spatial and temporal biases in sampling which may limit their use for tracking abundance. Often museum records are the only historical data available, and Boyle and colleagues make long-term abundance estimates for the Eastern North American Monarch butterfly (*Danaus plexippus*) and its milkweed hostplant (*Asclepias* spp.) using 1,191 and 31,510 records from 1900-2016, respectively. They conclude that Monarch and milkweed abundance started to decline in the mid-20^th^ century, before the adoption of herbicide-resistant crops that are often blamed for losses of Monarch hostplants. Using the same data, I argue that the Monarch trend changes with the choice of taxa used to standardize Monarch records. The abundance trend after dividing Monarch records by butterfly (Rhopalocera) or Nymphalidae records, instead of by Lepidoptera as in Boyle et al. (2019), shows no mid-century peak corresponding to the milkweed trends. One reason the Monarch trend reported by Boyle and colleagues changes when standardized by other taxa is the declining proportion of butterflies within Lepidoptera records from a peak of 40% in the mid-20^th^ century to less than 10%. This reanalysis shows that changes over time within the taxa used to standardize records matter, in addition to potential sampling biases in the species of interest.

---

Museum records can document long-term changes in phenology, species interactions, and trait evolution (1). However, these data have spatial and temporal biases in sampling which may limit their use for tracking abundance (2). Often museum records are the only historical data available, and Boyle and colleagues make long-term abundance estimates for the Eastern North American Monarch butterfly (*Danaus plexippus*) and its milkweed hostplant (*Asclepias* spp.) using 1,191 and 31,510 records from 1900-2016, respectively (3). They conclude that Monarch and milkweed abundance started to decline in the mid-20^th^ century, before the adoption of herbicide-resistant crops that are often blamed for losses of Monarch hostplants (4). Using the same data, I argue that the Monarch trend is sensitive to the method of standardization and appears less robust than the milkweed trend.

Boyle and colleagues recognize that museum records must be standardized by collection effort to estimate an index of annual relative abundance (2, 3, 5). They divide the number of Monarch records by the number of Lepidoptera records in each year. Their abundance index peaks mid-20^th^ century before a long-term decline (reproduced in the top row of Figure 1A). However, this trend changes with the choice of taxa used to standardize Monarch records. The abundance trend after dividing Monarch records by butterfly (Rhopalocera) or Nymphalidae records shows no mid-century peak corresponding to the milkweed trends (Figure 1A). I also show similar results from generalized linear models with linear and quadratic effects of year that account for the annual number of museum records with weights (5), a feature which the approach in (3) lacks (Figure 1B).

**Figure 1:**
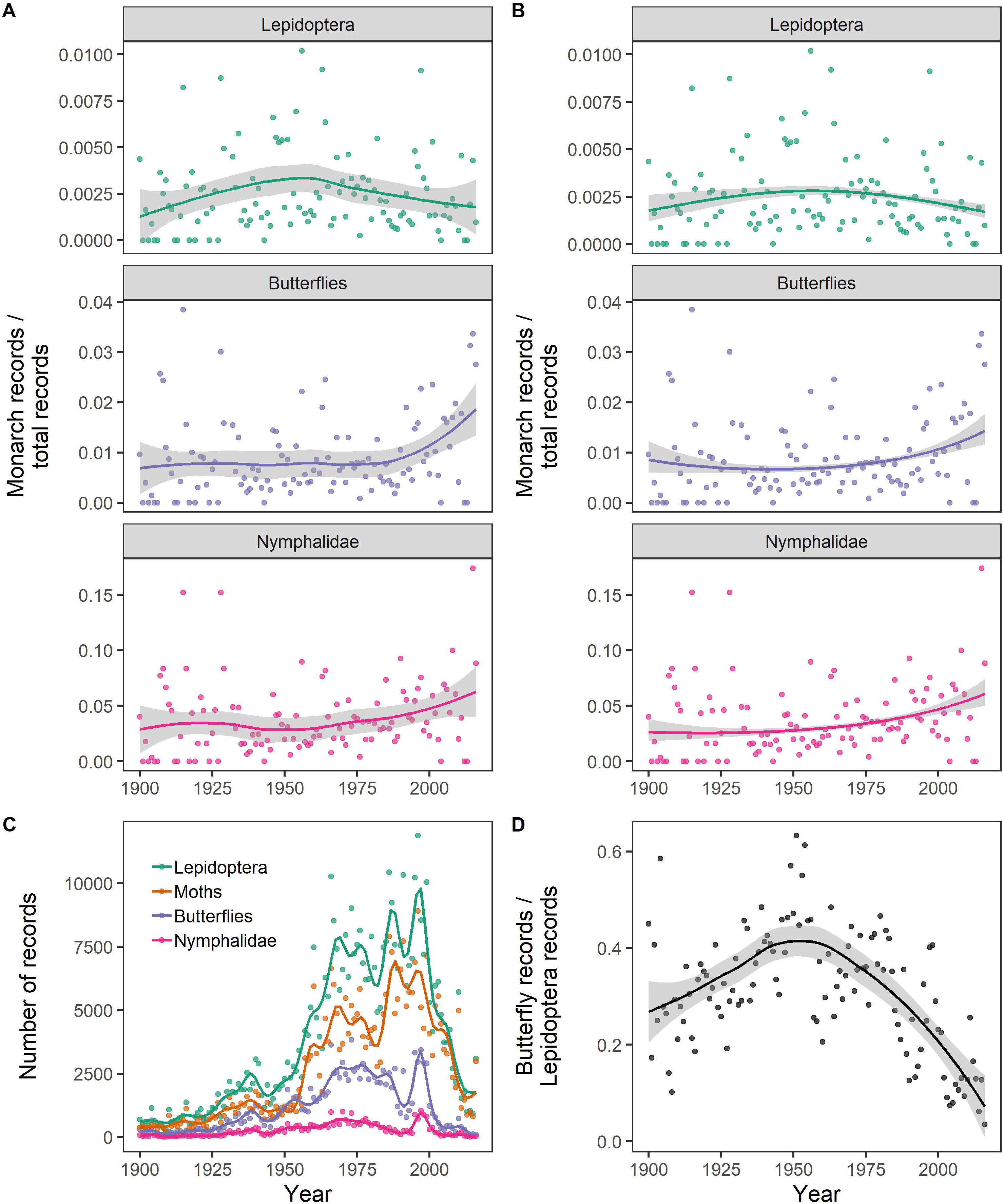
Trends in Eastern North American Monarch butterfly museum records change with the choice of standardization. All data came from (8) and span 1900-2015 and the Eastern USA. **A.** I reproduce Figure 1A in (3) with their standardization by Lepidoptera records and present two alternative standardizations (Rhopalocera and Nymphalidae). I similarly use the default LOESS smooth in the *ggplot2* R package for visualizing trends and 95% confidence intervals (9). **B.** The relative abundance of the three standardizations are alternatively modeled with a binomial generalized linear model, weighted by the annual number of records, predicting relative abundance with linear and quadratic year covariates. **C.** Total number of records of Lepidoptera, moths, butterflies, and Nymphalidae each year with splines showing trends. **D.** The proportion of butterfly records to all Lepidoptera records shows a strong temporal trend that influences the mid-20^th^ century peak of Monarch abundance reported in (3) and shown in the top row of **A** and **B**.

Collection effort that does not target the species of interest should be excluded when possible in these standardizations. Within the Lepidoptera, moths and butterflies would be most frequently sampled by nighttime light traps and daytime netting, respectively. One reason the Monarch trend reported in (3) changes when standardized by other taxa is the declining proportion of butterflies within Lepidoptera from a peak of 40% to less than 10% (Figures 1C & 1D), potentially due to increasing use of light traps around the mid-20^th^ century (6). In reference to museum records, Boyle and colleagues note that “the most concerning possible biases are those that change over time within a species” (3). This reanalysis shows that changes over time within the taxa used to standardize records also matter.

I do not think that this reanalysis presents the true Monarch trend, since it contrasts with recent declines (7). Rather, I think analysis of abundance from biological records needs more data and methodological advances to approach the value of systematic monitoring (2). The estimates for milkweed trends may be more robust with thirty times the number of herbarium records compared to Monarch specimens (3). Boyle and colleagues verify their method for herbarium records by correctly estimating increasing trends in four invasive plants over the 20^th^ century. A similar approach with invasive insects would be a valuable test to verify if museum records can estimate long-term trends in highly variable insect populations.

